# RMetD2: a tool for integration of relative transcriptomics data into Genome-scale metabolic models

**DOI:** 10.1101/663096

**Authors:** Cheng Zhang, Sunjae Lee, Gholamreza Bidkhori, Rui Benfeitas, Alen Lovric, Shuqi Chen, Mathias Uhlen, Jens Nielsen, Adil Mardinoglu

**Affiliations:** Science for Life Laboratory, KTH – Royal Institute of Technology, SE-171 21 Stockholm, Sweden; Department of Biology and Biological Engineering, Chalmers University of Technology, SE-412 96 Gothenburg, Sweden; Centre for Host–Microbiome Interactions, Dental Institute, King’s College London, London, SE1 9RT, United Kingdom

**Author notes:** **Correspondence:** Dr. Cheng Zhang, Science for Life Laboratory, KTH – Royal Institute of Technology, Tomtebodavägen 23A, SE-17165 and Stockholm, Sweden; Prof. Adil Mardinoglu, Science for Life Laboratory, KTH – Royal Institute of Technology, Tomtebodavägen 23A, SE-17165 and Stockholm, Sweden. Centre for Host–Microbiome Interactions, Dental Institute, King’s College London, London, SE1 9RT, United Kingdom (Sunjae Lee, Gholamreza Bidkhori).

**Keywords:** differential expression, genome-scale metabolic models, transcriptomics, flux balance analysis, systems biology

## Abstract

Relative Metabolic Differences version 2 (RMetD2) is a tool for integration of differentially expressed (DE) genes into genome-scale metabolic models (GEMs) for revealing the altered metabolism between two biological conditions. This method provides a robust evaluation of the metabolism by using flux ranges instead of a single set of flux distributions. RMetD2 classifies reactions into three different groups, namely up-regulated, down-regulated and unchanged, which enables systematic interpretation of the metabolic differences between two different conditions. We employed this method in three different case studies using mice and human datasets, and compared it with state-of-the-art methods used for studying condition-specific metabolic differences using GEMs. We observed that RMetD2 is capable of capturing experimentally-observed features that are missed by other methods, highlighting its potential use in biotechnology and systems medicine applications. RMetD2 is implemented in Matlab and it is available without any limitation at https://sourceforge.net/projects/rmetd.

## 1 Introduction

Genome-scale metabolic models (GEMs) are the comprehensive collections of known biochemical reactions and associated protein coding genes that are present in specific cells, tissues and organisms. They have been applied in metabolic engineering, antibiotic design, and increasingly used in revealing the underlying disease mechanism, discovery of biomarkers and identification of drug targets [1-7].

Many GEM based methods have been developed to help researchers understand biological data by integrating transcriptomic data obtained from different conditions [8]. Compared to simple enrichment analysis, GEM based methods could interpret the data in a connected manner based on network topology, and infer changes in remote pathways. For instances, Jerby et al. developed iMAT to construct context specific GEM for liver and predicted their hepatic flux changes which were later validated with experimental measurements [9]. In addition, Agren et al. developed a method called INIT which uses transcriptomic data and network topology of GEM to build cell type specific GEMs that exhibited different metabolic capabilities [10]. In another study, mCADRE was developed and used for reconstruction of GEMs for 126 human tissues and tumors, and proposed potential drug targets that specifically affect tumors based on their different topology [11].

Even though detailed insights can be gained based on network topology, predicting actual fluxes based on absolute transcriptomic data is a challenging task [12]. In this context, MADE [13], tFBA [14] and several other methods [15, 16] have been developed and applied to investigate the altered metabolism using GEMs and relative changes of gene expression. However, since there is a large solution space even when the objective function is optimized [17], it is difficult to identify consistently changed metabolic reactions using the above mentioned methods. On the other hand, there are also methods, such as reporter subnetworks [18], that identifies key metabolic subnetworks using relative transcriptomic data and the topological structure of GEMs. But these methods neglect stoichiometric information in the model and could likely miss some key reactions [19].

Here, we present the systematic formulation of a method called Relative Metabolic Differences version 2 (RMetD2). RMetD2 integrates relative gene expressions together with the stoichiometric structure of GEMs to identify key up-/down-regulated metabolic reactions that are most relevant to the changes in the transcriptome. The concept of RMetD has been successfully used in number of recent studies [4, 20, 21]. Unlike previous methods, RMetD2 stretches the potential biological effects into many steps by setting gradient constraints to the differentially expressed reactions. In this way, RMetD2 could be used to identify significantly up-/down-regulated reactions by correlation of flux ranges of all reactions, which allows users to prioritize and refine the key metabolic reactions and pathways, and enables detailed systematic interpretation of the biological data.

## 2 Materials and methods

### 2.1 GEM and simulation

Three GEMs have been employed for case studies. The first one is iHepatocyte2322, which is a GEM for human liver generated based on both transcriptomics and proteomics data in a previous study [22] and the irreversible format is used, which included 10565 reactions, 5195 metabolites and 2328 genes. The constraints for the model were taken from another study where the fluxes in liver tissue were measured and derived based on the body composition [4]. Mouse intestine GEMs, iMouse-Smallintestine, has been reconstructed in a previous study based on proteomics data, and the constraints for both conventional raised and germ-free mice are calculated based on the dietary consumption of the mice considering the effect of gut microbiota [4]. The irreversible version of this GEM has 8055 reactions, 4408 metabolites and 2337 genes. The secretion of chylomicron and HDL secretion were ensured by setting them as mandatory functions for the model according to the original study. The differentially expressed (DE) genes between the intestine of germ-free and conventionally raised mice were obtained from the same study [4]. The GEM for human cancer cell line HepG2 is generated in another recent study [21] based on transcriptomic data and exchange constraints calculated from the formulation of the cultivating medium. The irreversible form of the model which has 5403 reactions, 2784 metabolites and 1957 genes was used by RMetD2 in this study.

### 2.2 RMetD2

To clearly demonstrate the rationality of this method, we provide a toy network with 10 reactions and 3 genes as shown in Figure 1A. In this simplest case, if G1 which is associated with R3 is up-regulated, and G2 and G3 which are respectively associated with R5 and R8 are down-regulated, we could easily tell that the flux through R3 and R4 should be increased and fluxes via R10 should be decreased based on the topology. However, if G2 and G3 are changes in opposite direction, it will be tricky to predict the changes of flux via R10, since it will be dependent on the original flux scale and stoichiometric coefficients of both R6 and R9. And this scenario is very common in real cases studying biological changes in a genome-scale.

**Figure 1.**
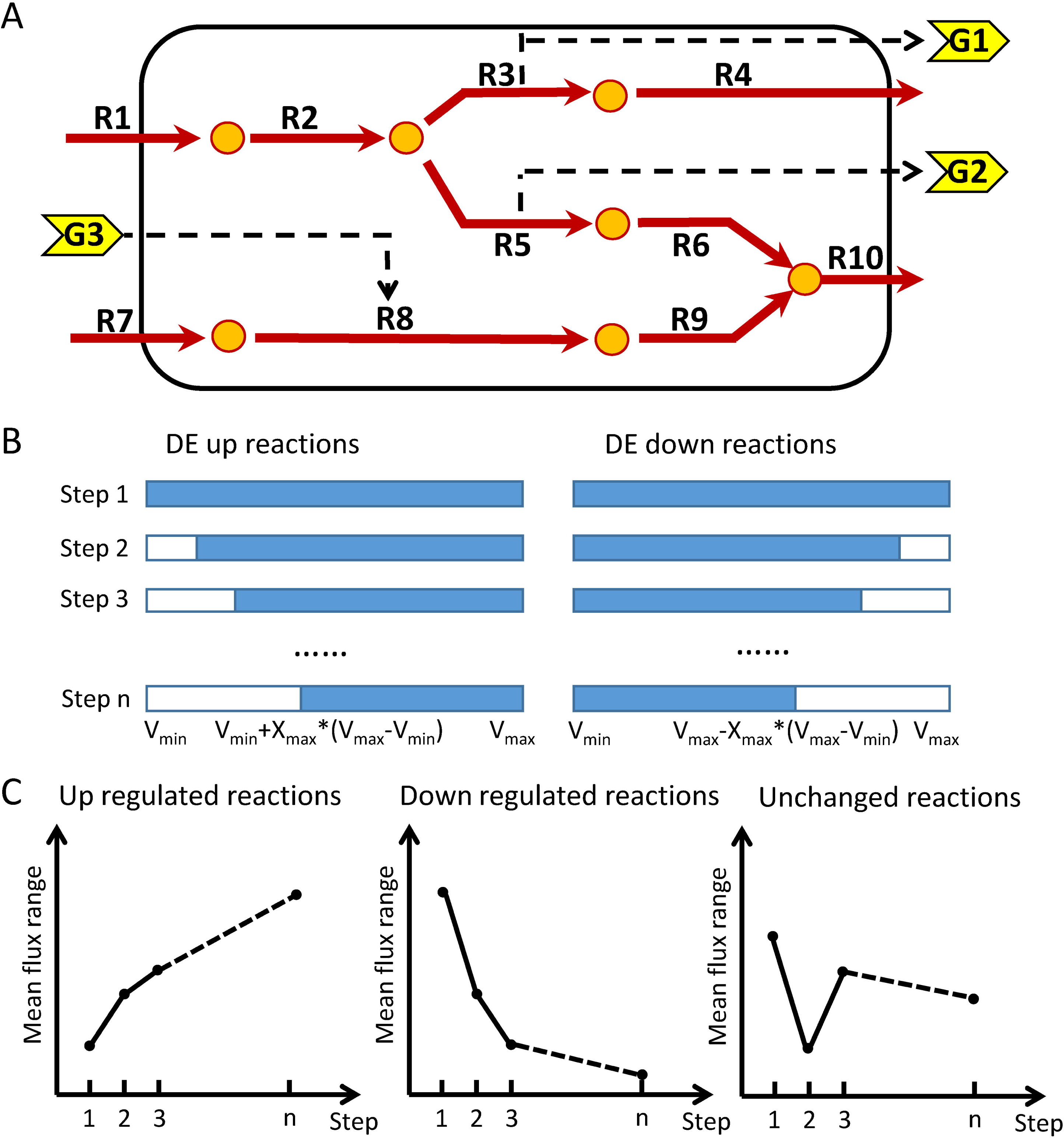
**B** (A) Toy model with solid red arrows, yellow circles, dashed arrows and yellow arrows representing metabolic reactions, metabolites, reaction-gene associations and metabolic genes, respectively. (B) Concept of constraining up- and down-regulated reactions in RMetD2. The constraints of lower bounds for reactions that are associated with up-regulated genes are pushed up step by step until they reach the maximal feasible limits, and vice versa. (C) Line plot examples showing the respective changes in flux ranges of up-, down-regulated and unchanged reactions. The x-axis indicates the average of the maximal and minimal flux range, and the y-axis indicates which step model it is.

The principal idea of RMetD2 is: to push the constraints of the flux for up- and down-regulated reactions respectively up and down using DE genes as indicator of directions (as shown in Figure 1B). The key concept of RMetD2 is to divide this process into several steps so that the consistency of the changes could be evaluated. The input for RMetD2 includes a GEM in either COBRA [23] or RAVEN [24] format (where objective function is optional), the DE genes and their fold changes (perturbed vs. reference). Additional constraints for the perturbed state model may optionally be added in case there are known metabolic differences between the two biological states (e.g. change of medium or gene knock-out). Otherwise, the original constraints of the input model are used. Using these inputs, RMetD2 classifies reactions into 3 different groups, namely up-regulated, down-regulated, and unchanged, based on the changes in flux ranges, and provide correlations and significance of relevance of each reaction to the biological change (as shown in Figure 1C). For instance, if G1 and G2 are up-regulated, and G3 is down-regulated, R4, R9 and R10 will probably be respectively classified as up-regulated, down-regulated and unchanged. It should be noted that, we used ‘constraints’ for the upper and lower bounds allowed for the reactions, and ‘flux ranges’ for the actual achievable maximum and minimal fluxes (with objective function maximized if there is any).

The implementation of RMetD2 includes several steps: (1) The given GEM is converted to an irreversible form so that all reactions will not carry any negative fluxes. (2) We identified up-/down-regulated reactions by associating DE genes using gene-protein-reaction relationships in GEMs. In brief, reactions associated with only up-(or down-) regulated genes are defined as up-(or down-) regulated genes. When a reaction is catalyzed by a protein complex that consists of several subunits and they show different expression change directionality, this reaction is defined as down-regulated since the down-regulation of one subunit will result in the decrease of the whole protein complex. In case of reactions catalyzed by multiple isozymes showing different expression directionalities, we define them as not significantly changed and leave it as it is in order to avoid false positives. (3) We calculated the upper and lower bounds for all up-/down-regulated reactions by flux variabilities analysis (FVA) based on the provided constraints (and if applicable, objective function), for both reference and perturbed model. (4) Then, we created *n* (default is 5) models in addition to the reference model and calculated flux ranges for all of the reactions by pushing the constraints to the same direction as the changes of related gene expression. The mathematical expression is defined as below:

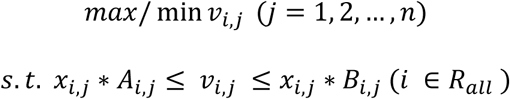

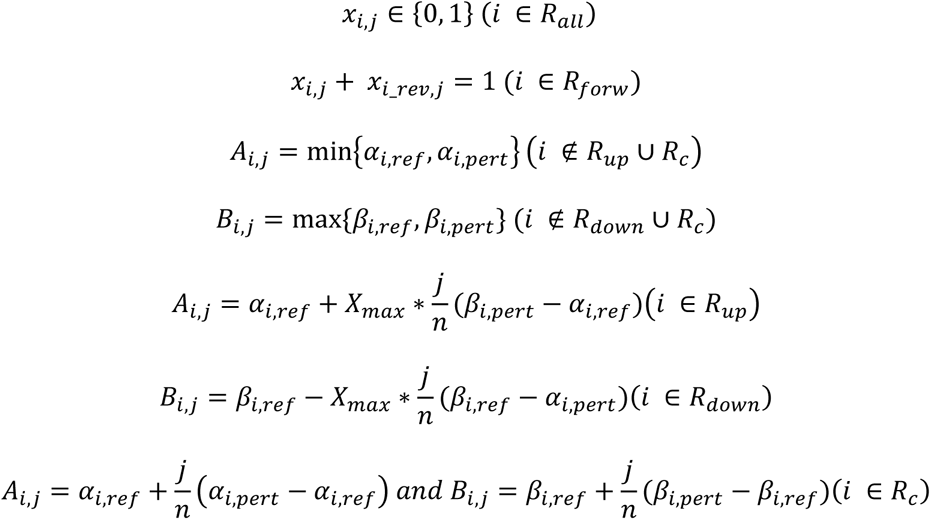

where *v*_*i,j*_ is the flux for *i*th reaction in *j*th model, *α*_*i,ref*_ and *β*_*i,ref*_ are upper and lower constraints for *i*th reaction in the reference model, whereas *α*_*i,pert*_ and *β*_*i,pert*_ are upper and lower constraints for *i*th reaction in the perturbed model. *R*_*all*_, *R*_*up*_, *R*_*down*_ and *R*_*c*_ represents all, up-regulated, down-regulated and constrained reaction sets, respectively. *R*_*forw*_ represents forward reactions of all reversible reactions in the original model. *x*_*i,j*_ is the binary variable defining whether *i*th reaction in *j*th model could carry flux, and *x*_*i_rev,j*_ is the binary variable of the reverse reaction of *i*th reaction. They are added to make sure that only one of the divided reversible reactions in irreversible model should carry flux and be constrained. *X*_*max*_ is a constant parameter calculated by bisectional searching, and introduced to push the fluxes as much into the same direction as the gene expression changes as possible but still in feasible ranges. (5) Finally, we defined every reaction as up-/down-regulated or unchanged based on the Spearman correlation between two vectors, *V*_*i*_and *X*, that calculated as below:

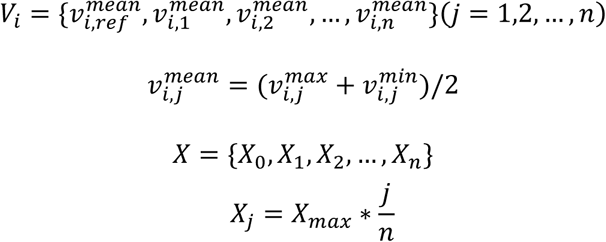

where *V*_*i*_ is the average flux vector of *i*th reaction in the model, and *X* is what we call step vector. Note that 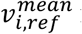 is the average of maximum and minimum flux range for *i*th reaction in reference model. If *V*_*i*_ is significantly positively/negatively correlated with *X* (P adjusted<0.05), the *i*th reaction is defined as up-/down-regulated, respectively; otherwise, it is defined as unchanged.

### 2.3 MADE

We compared the predictions of RMetD2 with those of MADE, a tool frequently used to compare two conditions using GEMs. MADE is implemented through the TIGER toolbox [25], and the time limit is set to 36000s. The flux ranges for all reactions were also calculated by the built-in function ‘fva’ in TIGER toolbox designed specifically for TIGER MADE GEMs.

### 2.4 Metabolic subsystem enrichment test

Each reaction in GEMs is annotated with a specific metabolic subsystem. Therefore, the significance of overlapping between reactions of different RMetD2 classes and metabolic subsystems of GEMs may be assessed using hypergeometric test. In this study, a RMetD2 class is enriched in a metabolic subsystem if the adjusted hypergeometric test P value of the overlap is less than 0.05 after Benjamini and Hockberg correction for multiple hypothesis testing.

For simple enrichment analysis, we used again GEMs for fair comparison. All genes associated with a specific metabolic subsystem in GEM is retrieved based on its gene-protein-reaction annotation, and up-/down-regulated genes are enriched in a metabolic system if the overlap is statistically significant using the same cutoff as metabolic subsystem enrichment test.

### 2.5 Availability and requirement

RMetD2 is a method build in Matlab (version R2017a) environment, and the source code is available at https://sourceforge.net/projects/rmetd/. In order to have fair comparison wherever relevant, the IBM ILOG CPLEX Optimization Studio v12.8 was employed for solving all linear programming and mixed integer linear programming in this study.

## 3 Results

### 3.1 Case 1: Liver before and after low-carbon diet

For benchmarking, a show case is provided where RMetD2 was used to reveal the metabolic differences in liver before and after carbon-restricted diet based on transcriptomics data generated in a previous study [4]. Tissue biopsy samples were taken from patients with fatty liver diseases before and one week after dietary change. In addition, the liver fat content is measured for both time points and found to be significantly decreased in subjects. The data used as an experimental benchmarking point for the *in silico* methods. We found that 952 and 3263 genes respectively significantly up- and down-regulated (FDR<0.05) between two different conditions. With implementation of RMetD2 with no predefined objective function, we identified 808 and 4987 significantly up- and down-regulated reactions (FDR<0.05), respectively. As expected, the reaction ‘HMR_9883’, which refers to triacylglycerol pool generation and thus indicates the accumulation of liver fat in the GEM is among the ones that are significantly down-regulated. For comparison, we also generated condition specific GEMs using MADE which is one of the most frequently used methods for comparing two conditions, and does not require predefined objective function. We compared the mean flux range before and after dietary change obtained by MADE. We identified 4267 and 811 reactions that respectively up- and down-regulated. Interestingly, as shown in Figure 2 we found that MADE predicted an increased flux through liver fat accumulation, which is the opposite to what we observed experimentally and computationally using RMetD2. Our analysis strongly demonstrated the advantage of using RMetD2 in biological applications.

**Figure 2.**
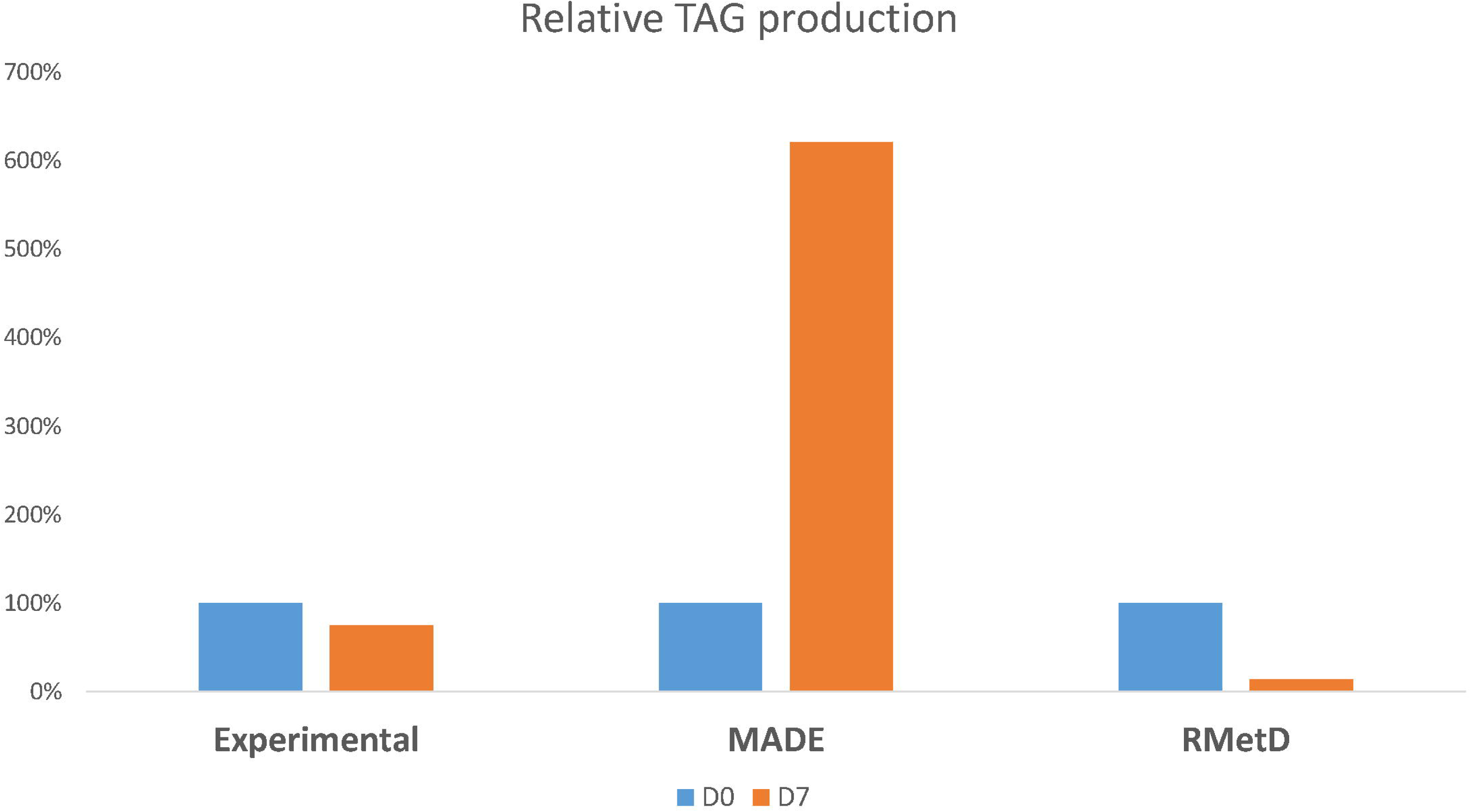
Barplot showing the relative liver TAG content at day 0 and day 7 for experimental measurement and in silico predictions generated by MADE and RMetD2. All values are normalized by the D0.

### 3.2 Case 2: Liver cancer cell line before and after gene inhibition

We also applied RMetD2 to investigate the metabolic difference between wild-type and PKLR inhibited liver cancer cell line, HepG2, using constraints and DE genes from our recent study [21]. The reference GEM of HepG2 was reconstructed in the same study based on transcriptomic data as well as composition of culture medium using tINIT [26]. The DE genes strongly related to silencing of PKLR (adjusted P <0.0001, n = 1242) were used as input for RMetD2. As a result, RMetD2 suggested 274 and 2078 reactions to be up- and down-regulated, respectively. Among the down-regulated reactions, we found ‘HCC_biomass’ and ‘TAG_accumulation’, and these suggest decreased growth and triacylglycerol accumulation in PKLR inhibited HepG2 cell line which was also validated experimentally in the same study. In addition, we also found ‘HMR_4381’, which is the conversion reaction from glucose-6-phosphate to fructose-6-phosphate and a key step in glycolysis, is classified as down-regulated. This suggested a decreased glucose consumption through glycolytic pathway, and as expected, we also found a deceased glucose consumption in PKLR inhibited cells in our experimental validation. These results further demonstrated the power of RMetD2.

### 3.3 Case 3: Intestine of conventional raised mice vs. germ-free mice

To further demonstrate the use of RMetD2 in integration of additional constraints, another show case is provided. RMetD2 was used to reveal the metabolic differences between germ-free and conventional raised mice based on the dietary inputs and DE genes obtained from the previous study [21]. Since most of the frequently used methods could not be used for integrating additional constraints together with transcriptomic changes, we only compared the RMetD2 result with experimental observations. In summary, we found that 323 and 616 genes significantly up- and down-regulated, respectively (FDR<0.05), and found that 130 and 2446 reactions are up- and down-regulated, respectively between between germ-free and conventional raised mice. In addition, we found that the maximal production of chylomicron in the last step model is decreased in conventional raised mice which is consistent with the conclusion of the original study [23]. Moreover, the HDL production reaction is among the down-regulated reactions, which is also in agreement with the previous study [23]. Additionally, the flux ranges for these two reactions are calculated with only constraint changes while neglecting the gene expression changes. The flux range of chylomicron secretion is decreased which is the same as we found using RMetD2, but the HDL production capability is increased. This implies that the key to infer the decrease of chylomicron is the constraint changes, while the decrease of HDL secretion seems like a result of transcriptomic changes.

In addition, we performed metabolic subsystems enrichment analysis for up- and down-regulated reactions obtained from RMetD2. As shown in Figure 3, the enriched up-regulated metabolic pathways in conventionally raised mice include histidine metabolism, phenylalanine, tyrosine and tryptophan metabolism, branch amino acids metabolism and oxidative phosphorylation. These might reflect the difference in diet, since we observed that conventional mice ate around 20% more in general. On the other hand, the down regulated reactions are enriched in many different pathways, including almost all fatty acids metabolic process such as beta oxidation of fatty acids, fatty acids biosynthesis, carnitine shuttle and glutathione metabolism. The down-regulation of these pathways suggests that there might be less free fatty acids in blood of conventional raised might to deal with. We also compared the enriched subsystems with the ones suggested by simple DE gene enrichment analysis, and we found many differences as shown in Figure 3. Most of the significantly enriched metabolic pathways disappeared in simple gene enrichment analysis, which is expected since it will miss metabolic pathways that have no association with DE genes but directly or indirectly linked with those significant changed pathways through the topology. One good example is glutathione metabolism, which is indirectly linked to many different metabolic pathways and account for the cofactor balancing in the whole system. Glutathione level is shown to be decreased in conventional raised mice according to RMetD2, and this is in very good agreement with the original study in which the key finding is that gut microbiota modulates host glutathione metabolism. However, simple enrichment analysis completely missed it, which highlighted the advantage of RMetD2.

**Figure 3.**
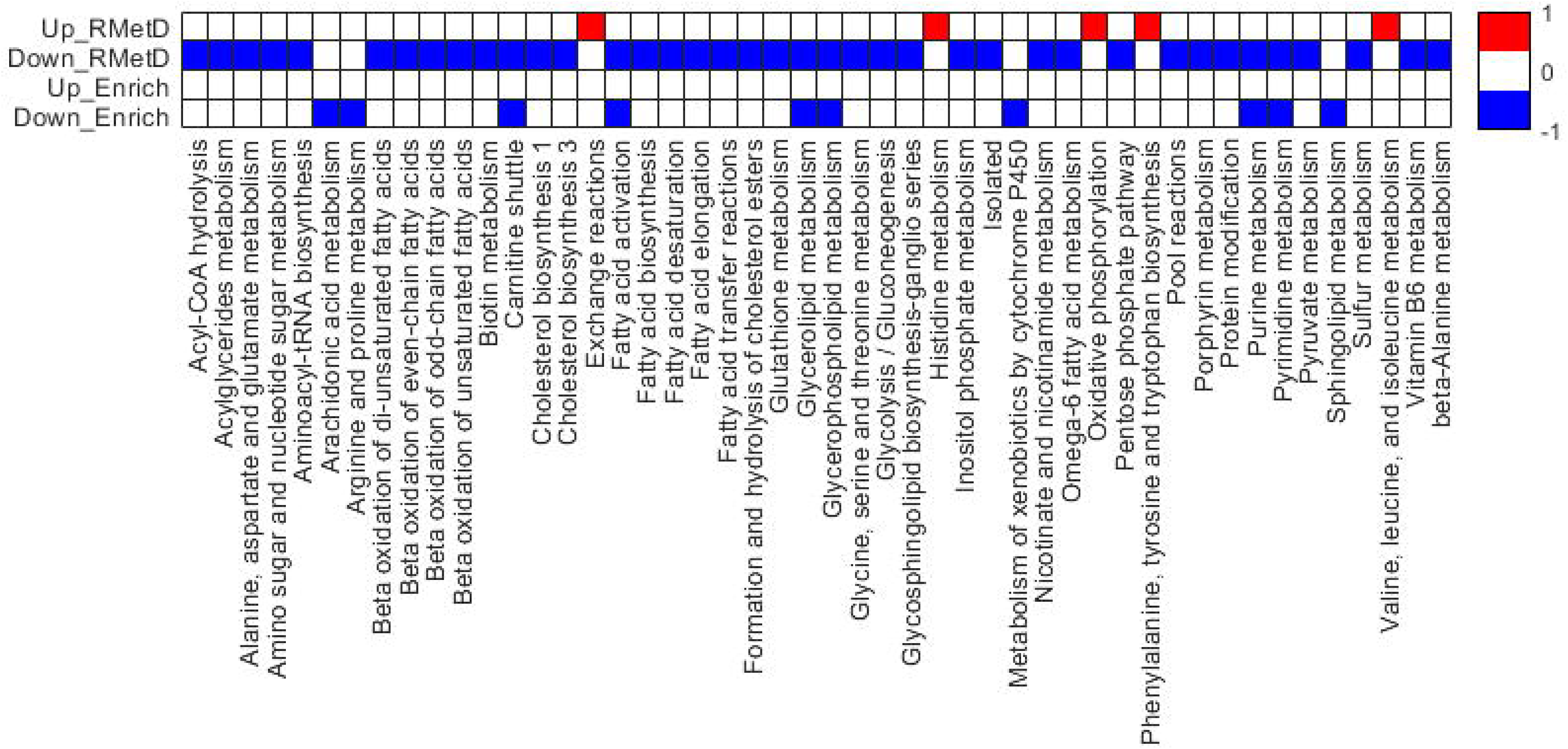
Heatmap showing the metabolic pathways enriched with up- and down-regulated reactions in intestinal of conventional raised mice compared to those of germ-free mice. The red and blue color in the heatmap represent significantly enriched with up-regulated and down-regulated reactions, respectively. The white color in the heatmap indicates there is no significant enrichment.

## 4 Discussion

In this study, we proposed a novel method called RMetD2 which can be used to integrate relative transcriptomic changes into GEMs and identify consistently up-/down-regulated reactions. We used RMetD for interpretation of the metabolic shift between two conditions at systems level. Unlike other methods, RMetD2 focuses on identifying relative change of reactions instead of having quantitative predictions since we are very well aware of the limitation of currently available data. To quantitatively predict the flux distribution using transcriptomics data, theoretically we need to know the translational efficiency of each gene, and kinetic values for all enzymes. Unfortunately, very few of them are currently available at genome-scale for any organism to the best of our knowledge. Therefore, we defined the main purpose of RMetD2 to identify key metabolic reactions or pathways in GEMs that are most relevant to the biological changes between conditions by integrating transcriptomic changes as well as any additional fluxomics data whenever it is available. In addition, RMetD2 could be implemented without predefined objective function, which is a great advantage for tissue modelling where there is no clear objective. We have shown in our case study that RMetD2 exhibited better predictions over MADE which is frequently used and thought to be among the best state-of-the-art methods.

It should be noted that in this method, gene-expression data are integrated as hard constraints which is the same as many other methods, such as GX-FBA [27]. Therefore, falsely annotated genes might have a significant impact on the results. So, the use of high quality GEM is very important and will likely improve the performance of RMetD2. Therefore, we are confident that the method presented here is promising and of great interest to researchers in the biotechnology as well as systems biology and medicine fields.

## Acknowledgement

This work was funded by the Knut and Alice Wallenberg Foundation.

## Author contribution

AM and CZ conducted the study. CZ developed the method. SJ, GR, SC and RB helped in development of the method. CZ performed most of the analysis and AL, SC, GR helped in the analysis. CZ wrote the paper and SJ, GR, RB, AL, SC, MU, JN, AM were involved in editing the paper.

## REFERENCES

[1] Mardinoglu, A., Boren, J., Smith, U., Uhlen, M., Nielsen, J., Systems biology in hepatology: approaches and applications. Nat Rev Gastroenterol Hepatol 2018.

[2] Turanli, B., Grøötli, M., Borén, J., Nielsen, J., et al., Drug repositioning for effective prostate cancer treatment. Frontiers in Physiology 2018, 9, 500.

[3] Benfeitas, R., Uhlen, M., Nielsen, J., Mardinoglu, A., New Challenges to Study Heterogeneity in Cancer Redox Metabolism. Front Cell Dev Biol 2017, 5, 65.

[4] Mardinoglu, A., Wu, H., Bjornson, E., Zhang, C., et al., An Integrated Understanding of the Rapid Metabolic Benefits of a Carbohydrate-Restricted Diet on Hepatic Steatosis in Humans. Cell Metab 2018, 27, 559–571 e555.

[5] Zhang, C., Hua, Q., Applications of Genome-Scale Metabolic Models in Biotechnology and Systems Medicine. Frontiers in Physiology 2016, 6, 413.

[6] O’Brien, E. J., Monk, J. M., Palsson, B. O., Using Genome-scale Models to Predict Biological Capabilities. Cell 2015, 161, 971–987.

[7] Mardinoglu, A., Nielsen, J., New paradigms for metabolic modeling of human cells. Curr Opin Biotechnol 2015, 34, 91–97.

[8] Blazier, A. S., Papin, J. A., Integration of expression data in genome-scale metabolic network reconstructions. Front Physiol 2012, 3, 299.

[9] Jerby, L., Shlomi, T., Ruppin, E., Computational reconstruction of tissue-specific metabolic models: application to human liver metabolism. Mol Syst Biol 2010, 6, 401.

[10] Agren, R., Bordel, S., Mardinoglu, A., Pornputtapong, N., et al., Reconstruction of Genome-Scale Active Metabolic Networks for 69 Human Cell Types and 16 Cancer Types Using INIT. PLoS Comput Biol 2012, 8.

[11] Wang, Y. L., Eddy, J. A., Price, N. D., Reconstruction of genome-scale metabolic models for 126 human tissues using mCADRE. Bmc Syst Biol 2012, 6.

[12] Machado, D., Herrgard, M., Systematic evaluation of methods for integration of transcriptomic data into constraint-based models of metabolism. PLoS Comput Biol 2014, 10, e1003580.

[13] Jensen, P. A., Papin, J. A., Functional integration of a metabolic network model and expression data without arbitrary thresholding. Bioinformatics 2011, 27, 541–547.

[14] van Berlo, R. J., de Ridder, D., Daran, J. M., Daran-Lapujade, P. A., et al., Predicting metabolic fluxes using gene expression differences as constraints. IEEE/ACM Trans Comput Biol Bioinform 2011, 8, 206–216.

[15] Fang, X., Wallqvist, A., Reifman, J., Modeling phenotypic metabolic adaptations of Mycobacterium tuberculosis H37Rv under hypoxia. PLoS Comput Biol 2012, 8, e1002688.

[16] Navid, A., Almaas, E., Genome-level transcription data of Yersinia pestis analyzed with a new metabolic constraint-based approach. Bmc Syst Biol 2012, 6, 150.

[17] Reed, J. L., Shrinking the metabolic solution space using experimental datasets. PLoS Comput Biol 2012, 8, e1002662.

[18] Patil, K. R., Nielsen, J., Uncovering transcriptional regulation of metabolism by using metabolic network topology. P Natl Acad Sci USA 2005, 102, 2685–2689.

[19] Zhang, C., Hua, Q., Applications of Genome-Scale Metabolic Models in Biotechnology and Systems Medicine. Front Physiol 2015, 6, 413.

[20] Mardinoglu, A., Shoaie, S., Bergentall, M., Ghaffari, P., et al., The gut microbiota modulates host amino acid and glutathione metabolism in mice. Mol Syst Biol 2015, 11.

[21] Liu, Z., Zhang, C., Lee, S., Kim, W., et al., Pyruvate kinase L/R is a regulator of lipid metabolism and mitochondrial function. Metabolic Engineering 2019, 52, 263–272.

[22] Mardinoglu, A., Agren, R., Kampf, C., Asplund, A., et al., Genome-scale metabolic modelling of hepatocytes reveals serine deficiency in patients with non-alcoholic fatty liver disease. Nat Commun 2014, 5.

[23] Schellenberger, J., Que, R., Fleming, R. M., Thiele, I., et al., Quantitative prediction of cellular metabolism with constraint-based models: the COBRA Toolbox v2.0. Nat Protoc 2011, 6, 1290–1307.

[24] Agren, R., Liu, L., Shoaie, S., Vongsangnak, W., et al., The RAVEN toolbox and its use for generating a genome-scale metabolic model for Penicillium chrysogenum. PLoS Comput Biol 2013, 9, e1002980.

[25] Jensen, P. A., Lutz, K. A., Papin, J. A., TIGER: Toolbox for integrating genome-scale metabolic models, expression data, and transcriptional regulatory networks. BMC Syst Biol 2011, 5, 147.

[26] Agren, R., Mardinoglu, A., Asplund, A., Kampf, C., et al., Identification of anticancer drugs for hepatocellular carcinoma through personalized genome-scale metabolic modeling. Mol Syst Biol 2014, 10, 721.

[27] Navid, A., Almaas, E., Genome-level transcription data of Yersinia pestis analyzed with a New metabolic constraint-based approach. Bmc Syst Biol 2012, 6.

